# Development of an improved inhibitor of Lats kinases to promote regeneration of mammalian organs

**DOI:** 10.1101/2022.04.07.487521

**Authors:** Nathaniel R. Kastan, Sanyukta Oak, Rui Liang, Leigh Baxt, Robert W. Myers, John Ginn, Nigel Liverton, David J. Huggins, John Pichardo, Matthew Paul, Thomas S. Carroll, Aaron Nagiel, Ksenia Gnedeva, A. J. Hudspeth

**Affiliations:** Howard Hughes Medical Institute and Laboratory of Sensory Neuroscience, The Rockefeller University, New York, NY 10065; Tri-Institutional Therapeutics Discovery Institute, New York, NY 10021, USA; Department of Physiology and Biophysics, Weill Cornell Medical College of Cornell University, New York, NY 10065, USA; Bioinformatics Resource Center, The Rockefeller University, New York, NY 10065; The Vision Center, Department of Surgery, Children’s Hospital Los Angeles, Los Angeles, CA, USA; The Saban Research Institute, Children’s Hospital Los Angeles, Los Angeles, CA, USA; Roski Eye Institute, Department of Ophthalmology, Keck School of Medicine, University of Southern California, Los Angeles, CA, USA; Eli and Edythe Broad CIRM Center for Regenerative Medicine and Stem Cell Research, University of Southern California, Los Angeles, CA 90033, USA; Tina and Rick Caruso Department of Otolaryngology—Head and Neck Surgery, University of Southern California, Los Angeles, CA 90033, USA

**Keywords:** cardiomyocyte, HEK293A cell, mouse, regeneration, retinal organoid, TRULI

## Abstract

The Hippo signaling pathway acts as a brake on regeneration in many tissues. This cascade of kinases culminates in the phosphorylation of the transcriptional cofactors Yap and Taz, whose concentration in the nucleus consequently remains low. Various types of cellular stress can reduce phosphorylation, however, resulting in the accumulation of Yap and Taz in the nucleus and subsequently in mitosis. We earlier identified a small molecule, TRULI, that blocks the final kinases in the pathway, Lats1 and Lats2, and thus elicits proliferation of several cell types that are ordinarily post-mitotic and aids regeneration in mammals. In the present study we present the results of chemical modification of the original compound and demonstrate that a derivative, TDI-011536, is an effective blocker of Lats kinases *in vitro* at nanomolar concentrations. The compound fosters extensive proliferation in retinal organoids derived from human induced pluripotent stem cells. Intraperitoneal administration of the substance to mice suppresses Yap phosphorylation for several hours and induces transcriptional activation of its target genes in the heart, liver, and skin. Moreover, the compound initiates the proliferation of cardiomyocytes in adult mice following cardiac cryolesions. After further chemical refinement, related compounds might prove useful in protective and regenerative therapies.

**Significance Statement:** In humans and other mammals, many organs regenerate through the proliferation of cells that replace those that have succumbed to aging or injury. However, proliferation is largely absent in certain critical organs, including the heart, the central nervous system, and sensory organs such as the inner ear and retina. The Hippo-Yap biochemical signaling pathway, a cascade of proteins that—when active—inhibits cell division, constitutes one impediment to proliferation. We earlier identified a small molecule that interrupts Hippo-Yap signaling and thus relieves this block for some non-proliferating cells *in vitro*. In the present investigation, we have chemically modified the original substance to yield a more potent analog that is effective for several hours in mammalian tissues *in vivo* and initiates the proliferation of heart-muscle cells after cryolesioning. After further refinements, compounds of this family might prove useful in regenerative therapies.

## Introduction

The highly conserved Hippo pathway is a key participant in numerous physiological processes and pathologies, including development, homeostasis, regeneration, cancer, and neurodegeneration (1–4). Hippo signaling integrates information from the cellular environment, including biomechanical cues, cell density, cell polarity, metabolic challenges, and signals such as Notch and Wnt. The core pathway comprises two pairs of core kinases. First, when activated by upstream signals, Mst1 and Mst2 phosphorylate Lats1 and Lats2. The latter kinases then phosphorylate the transcriptional co-activators Yap and Taz, which largely remain in the cytoplasm. When the kinase cascade is inactive, however, unphosphorylated Yap accumulates in the nucleus, where it interacts with transcription factors of the Tead family to initiate cell division (5, 6)(7).

Although transgenic approaches have elucidated much about the Hippo pathway, small molecules targeting the pathway offer a simplified avenue of investigation and could potentially have therapeutic utility. Over the last few years, a few compounds that target the Hippo pathway have been characterized. XMU-MP1, an inhibitor of the MST kinases, augments the regeneration of organs in which the process occurs naturally, such as the liver and intestine (8). Quinolinols, which promote Yap-dependent transcription by stabilizing the interaction between Yap and Teads, enhance the healing of skin wounds in mice (9). An additional Lats inhibitor, GA-017, has recently been shown to augment the creation of mouse intestinal organoids *in vitro* (10).

In the hope of identifying substances that promote proliferation in supporting cells of the internal ear, we earlier screened small molecules for compounds that facilitate the nuclear enrichment of Yap. From this effort emerged an inhibitor of Lats kinases, TRULI, which proved effective not only for its original objective, but also for promoting the proliferation of inner-ear sensory epithelia, cultured primary cardiomyocytes, and human retinal organoids (11). We have subsequently conducted a medicinal-chemistry campaign to enhance the potency and drug-like properties of Lats inhibitors of this family and to improve their efficacy *in vivo*. Here we characterize TDI-011536, a derivative that demonstrates a marked improvement in potency and physicochemical properties and offers a simple paradigm for systemic Lats inhibition in several mammalian tissues *in vitro* and *in vivo*.

## Results

### Enhancement of TRULI’s activity through medicinal chemistry

The original lead inhibitor of Lats kinases, TRULI, comprises a thiazolidine-2-imine core to which a hydrophobic benzyl moiety is linked through the ring’s nitrogen atom. In addition, a 7-azaindole-3-carboxylate substituent is attached to the core at the imine nitrogen (Fig. 1A). Although TRULI demonstrated good biochemical potency, its cellular potency was modest and it exhibited poor stability on exposure to mouse liver microsomes. With the aim of identifying compounds of greater potency and better drug-like properties, we systematically modified TRULI. To assess the potency of the new analogs, we measured their activity in the *in vitro* kinase assay and a cellular assay of Yap phosphorylation (11). The ATP concentration was increased from 10 µM to 2 mM because the increased potency of the new compounds, which were predicted to act as an ATP-competitive inhibitors of Lats kinases, prevented their evaluation in the original assay format. In addition, a value of 2 mM approximates the physiologically relevant ATP concentration.

**Figure 1.**
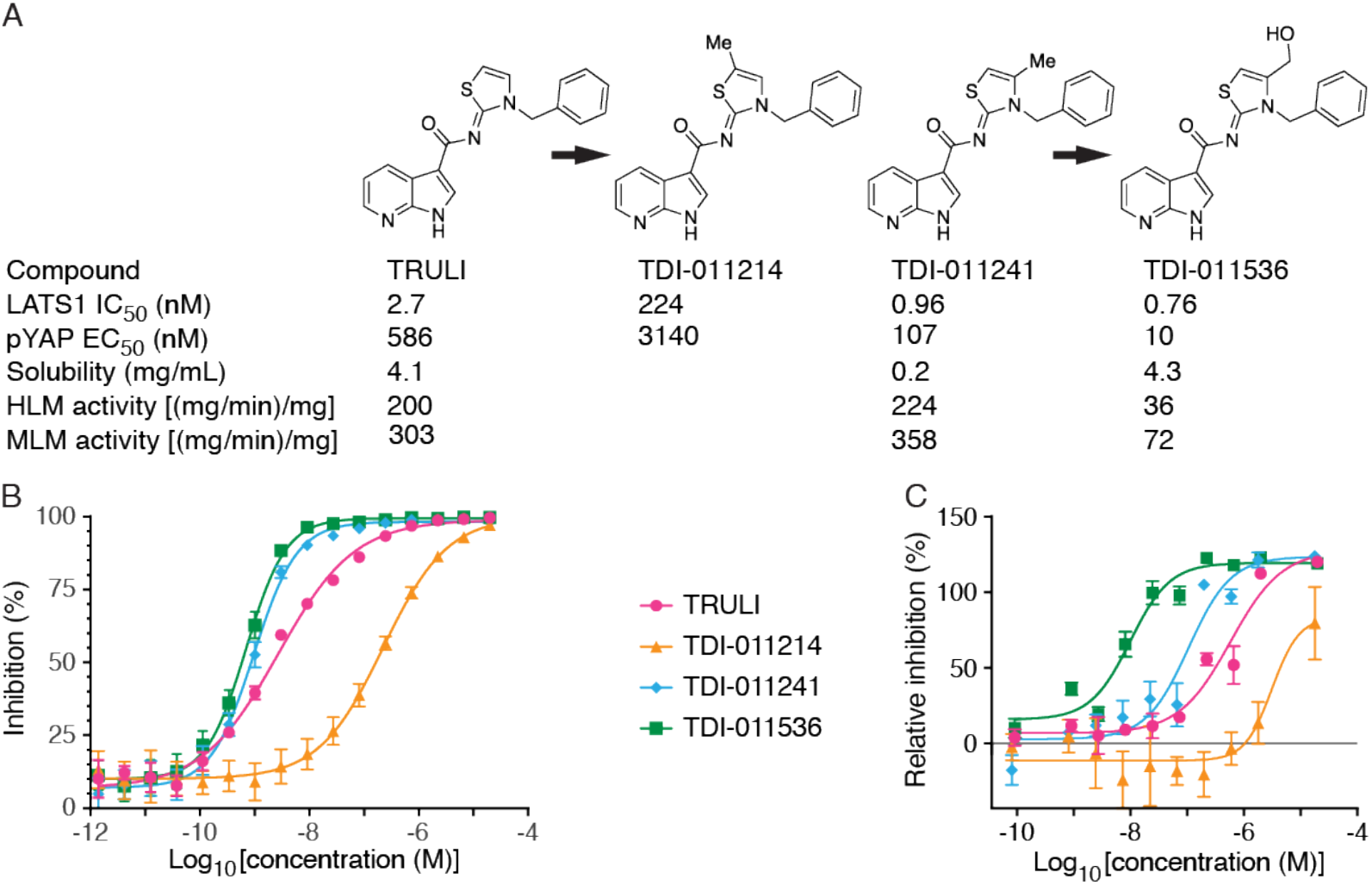
Development of an effective Lats kinase inhibitor. (*A*) The enhancement of the original Lats kinase inhibitor involved derivatization of the thiazolidine ring with a methyl and then a hydroxymethyl group. *IC*_50_ determinations were conducted in the presence of 2 mM ATP; aqueous solubility was measured at pH 6.8. HLM, human liver microsome; MLM, mouse liver microsome. (*B*) By an *in vitro* assay of Lats inhibition, methylation of the thiazolidine ring adjacent to the sulfur atom (TDI-011214) greatly diminished potency with respect to the original inhibitor TRULI. Methylation adjacent to the ring’s nitrogen atom (TDI-011241) enhanced potency, and the hydroxymethyl derivative TDI-011536 was more effective still. Error bars denote standard errors of the means (*n* = 4 measurements apiece in *N* = 4 independent experiments). (*C*) The relative inhibition of Lats kinases was determined from the ratio of pYap to tYap in homogenates from treated HEK293A cells. Although the order of efficacy of the four compounds was identical, the performance of TDI-011536 was strikingly superior. Error bars denote standard errors of the means (*n* = 2 measurements apiece in *N* = 2 independent experiments). For a total of 30 measurements for TDI-011536, *EC*_50_ = 40.8 ± 4.5 nM (mean ± SEM).

Computational docking studies suggested that the azaindole moiety, like that of ATP, binds to the hinge region of Lats kinases (11). Initial exploration of structure-activity relationships revealed limited opportunities to modify this region. We therefore retained the azaindole hinge-binding element and focused on the thiazolidine core. Examination of the docking model of TRULI in Lats1 suggested that there was space to extend deeper into the pocket located beneath the thiazolidine-2*-*imine core. Exploration of this hypothesis with simple methyl substitution provided the 5- and 4-methyl substituted compounds TDI-011214 and TDI-011241, respectively (Fig. 1A). The 5-methyl substitution in the former compound potentially oriented the lipophilic methyl group toward a polar region. In line with this possibility, TDI-11214 suffered a significant loss of enzymatic potency. However, methylation in the adjacent 4 position yielded improvements in both biochemical and cellular potency.

Despite the improved activity of TDI-011241, the increased lipophilicity resulted in poor aqueous solubility at pH 6.8; moreover, the compound also suffered from low metabolic stability (Fig. 1A). We sought to improve those physical properties by incorporating polar substituents. Enhancing the 4-methyl with a hydroxyl group yielded TDI-011536, which gratifyingly offered a significant gain in cellular potency as well as improved aqueous solubility and metabolic stability. Examination of TDI-011536 in our docking model suggests that the pendant hydroxyl group makes a hydrogen bond with Asp789. That the gain in cellular potency is not reflected in changes in biochemical activity suggests that the current assay is at the lower limit of detection based on the protein concentration of 555 pM.

### Effect of TDI-011536 on human retinal organoids

We have previously demonstrated that TRULI can decrease the phosphorylation of Yap and cause the proliferation of Müller glia in human retinal organoids *in vitro* (11). To test the potency of TDI-011536 in the same system, we applied the two compounds to retinal organoids derived from human induced pluripotent stem cells. After 24 hours of Lats kinase inhibition with 10 µM TRULI, there was a significant 30 % reduction in Yap phosphorylation as determined by the ratio of phospho-Yap (pYap) to total Yap (tYap; Fig. 2A,B). After treatment with 3 µM TDI-011536, Yap phosphorylation was reduced by almost 80 % from the level in containing an identical amount of dimethyl sulfoxide (DMSO), the vehicle used to solubilize the compound. We next tested the effect of increased Yap signaling on proliferation in Müller glia, which we assessed in organoids cultured in the presence of the thymidine analog 5-ethynyl-2′-deoxyuridine (EdU). After a five-day treatment with 10 µM TRULI, there was a robust, almost fivefold increase in the number of Sox9-positive Müller glia that incorporated EdU (Fig. 2C,D). By contrast, TDI-011536 treatment increased the number of doubly positive cells tenfold. These data confirm that inhibition of Lats kinases stimulates Yap signaling and the proliferation of Müller glia in human retinal organoids and demonstrate that TDI-011536 is significantly more potent than TRULI in this system. In conjunction with the improved solubility and metabolic stability of the compound, the results advanced TDI-11536 as an appropriate tool compound for investigating LATS inhibition *in vivo*.

**Figure 2.**
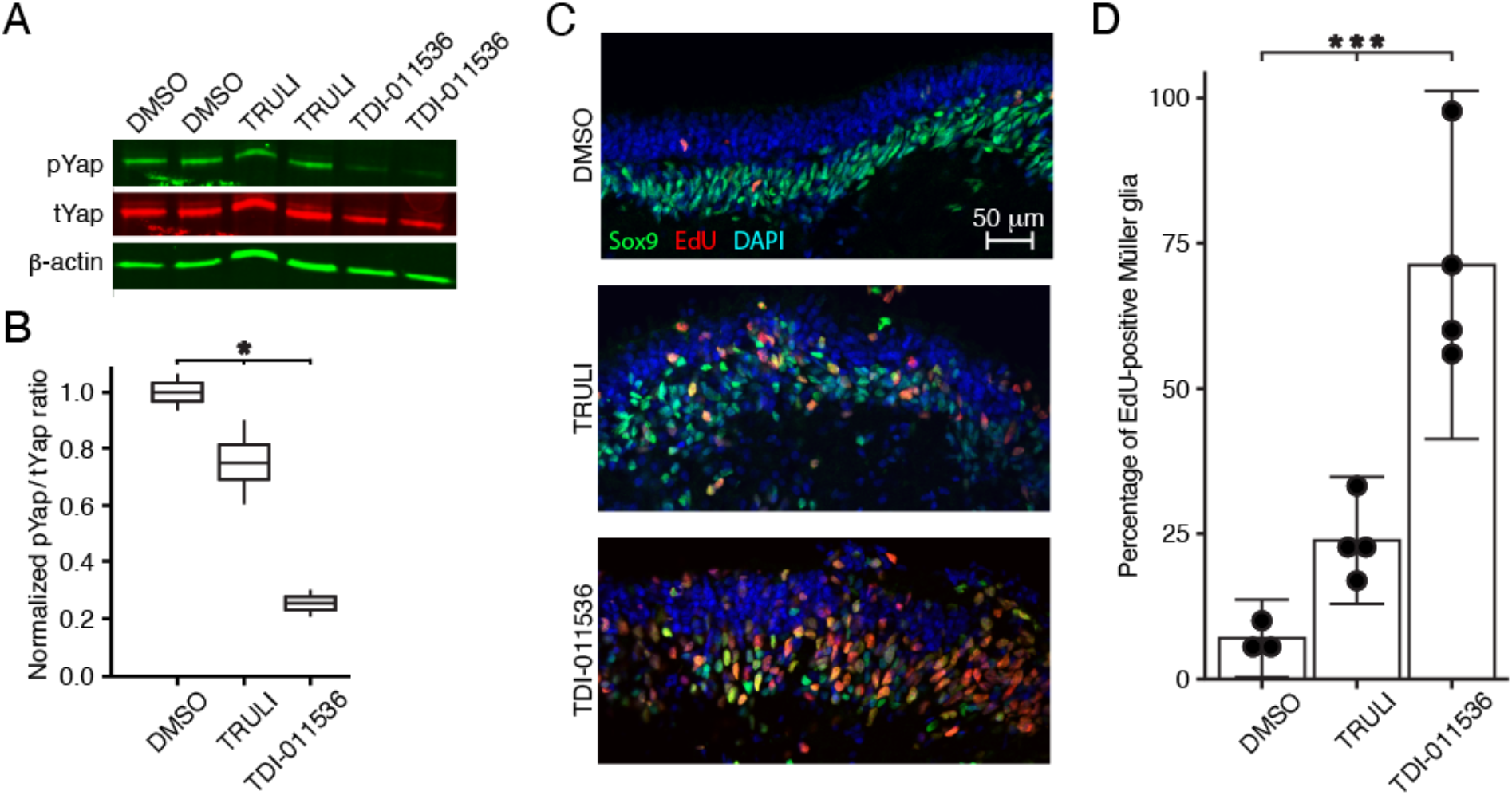
Proliferation of Müller glia in human retinal organoids after treatment with Lats kinase inhibitors. (*A*) An immunoblot for pYap and tYap after a 24 hr incubation of human retinal organoids in DMSO, 10 µM TRULI, or 3 µM TDI-011536 shows modest suppression of Yap phosphorylation after the former treatment and a profound effect of the latter. DMSO is the solvation vehicle for the compounds; β-actin serves as a loading control. (*B*) Quantification of the immunoblot by the ratio of pYap to tYap, both normalized to the DMSO control, confirms a significant effect of treatment with both compounds by one-way ANOVA, *P* = 0.019 for two experiments in each condition. (*C*) After five days of treatment, confocal immunofluorescence microscopy of sections from human retinal organoids reveals a substantial increase in cells doubly positive for EdU and Sox9, which represent proliferating Müller glia. (*D*) Quantification of the foregoing result demonstrates the significance of the effect by one-way ANOVA, *P* = 0.0003 for four experiments in each condition.

### *In vivo* activity of TDI-011536

To explore the efficacy of TDI-011536 in living animals, we conducted intraperitoneal injections of mice with a relatively high dose of 200 mg per kilogram of body weight (mg/kg). The compound was dissolved in Kolliphor HS 15, a vehicle that was also injected into control animals. Immunoblot analysis of the heart, liver, and skin immediately after injection demonstrated the high concentration of phosphorylated Yap (pYap) characteristic of control conditions. Two or four hours after injection, however, all three organs in treated animals demonstrated a remarkable reduction in pYap relative to total Yap (tYap; Fig. 3A). This process was reversible; by eight hours pYap levels returned to their baseline levels (SI Appendix, Fig. S1).

**Figure 3.**
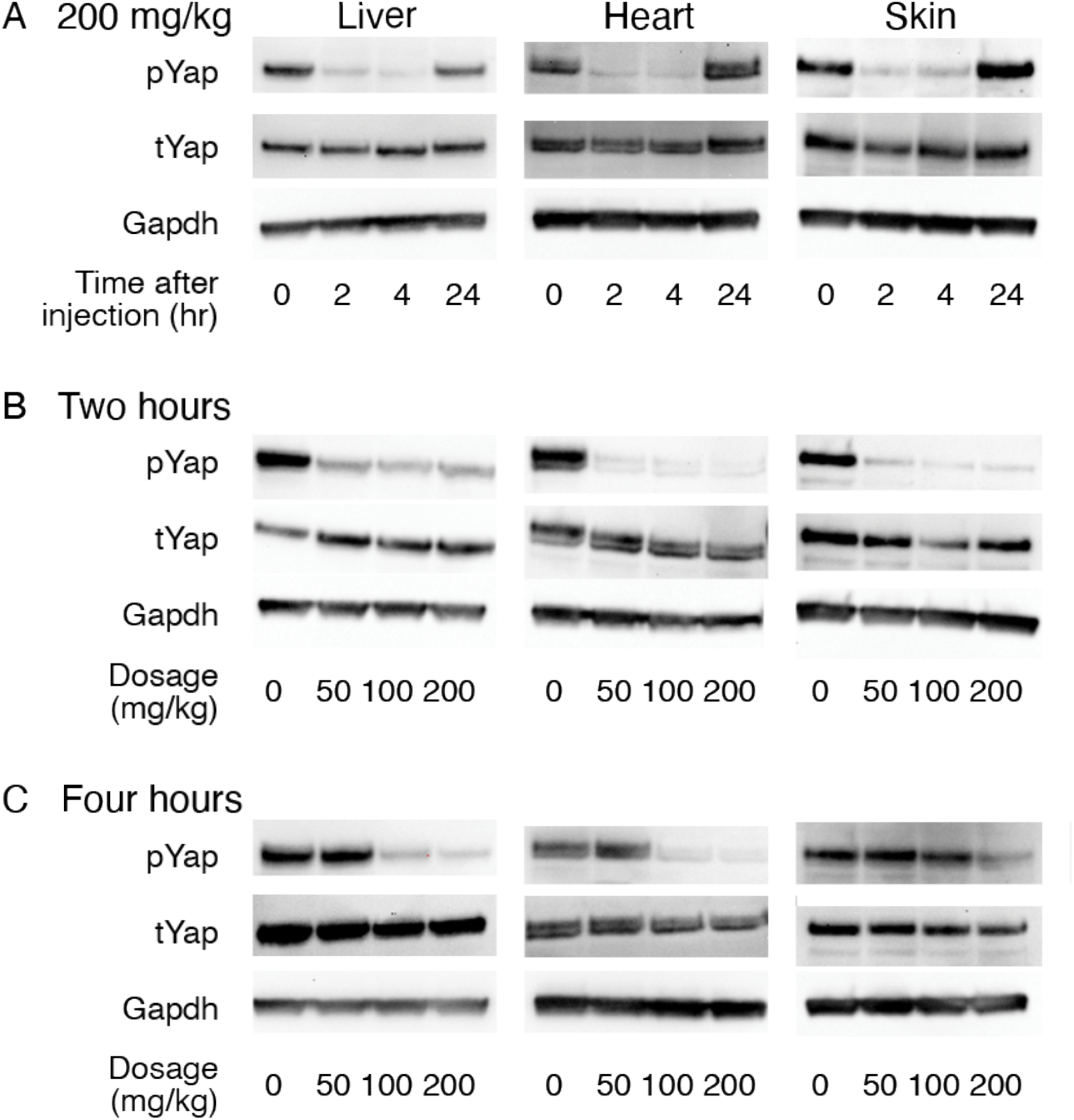
Engagement of target organs *in vivo*. (*A*) Immunoblots portray the amounts of Yap phosphorylated at residue S127 (pYap) and the total amounts of Yap (tYap) in the liver, heart, and skin. Injection of TDI-011536 at a dosage of 200 mg/kg greatly reduces the amount of pYap for at least four hours after injection in all three organs. The levels return to control values within a day. Glyceraldehyde 3-phosphate dehydrogenase (Gapdh) is included in each instance as a loading control. (*B*) Two hours after injection, all three dosages of TDI-011536 largely suppress the amount of pYap. (*C*) By four hours after injection, pYap has recovered following injections at 50 mg/kg. The concentration of pYap remains low in the heart and liver following injections at 100 mg/kg and 200 mg/kg, but largely recovers in the skin at the former dosage.

We next evaluated the effect of dosage de-escalation on the pharmacokinetics. We compared the original 200 mg/kg dose to injections of 100 mg/kg and 50 mg/kg. All three treatments evoked profound reductions of pYap in the three organs within two hours of injection (Fig. 3B). After four hours, however, the 50 mg/kg dose no longer diminished pYap significantly in any organ (Fig. 3C). The 100 mg/kg dose maintained the dephosphorylation of Yap through four hours in the liver and heart, but failed to do so in the skin. These data together demonstrate the concentration dependence of TDI-011536 activity *in vivo* and suggest that an injection of 200 mg/kg provides over four hours of Lats inhibition in the liver, heart, and skin.

### Effects of Lats kinase inhibition on gene expression

To confirm that treatment with TDI-011536 activates Yap *in vivo*, we collected livers and hearts for RNA sequencing four hours after injection at 200 mg/kg. Principal-component analysis of gene expression revealed that the samples clustered first by organ and second by treatment (Fig. 4A). We used gene set-enrichment analysis to examine changes in targets of Yap activity. In both the liver and heart, genes characteristically activated upon Yap expression displayed significant enrichment after exposure to TDI-011536 (Fig. 4B,C). Although some genes associated with the G1/S and G2/M phases of mitosis changed significantly in the liver, the set as a whole was not significantly enriched in either tissue (SI Appendix, Fig. S2A). This result might be expected after only a few hours of Yap activity, especially in the post-mitotic heart.

**Figure 4.**
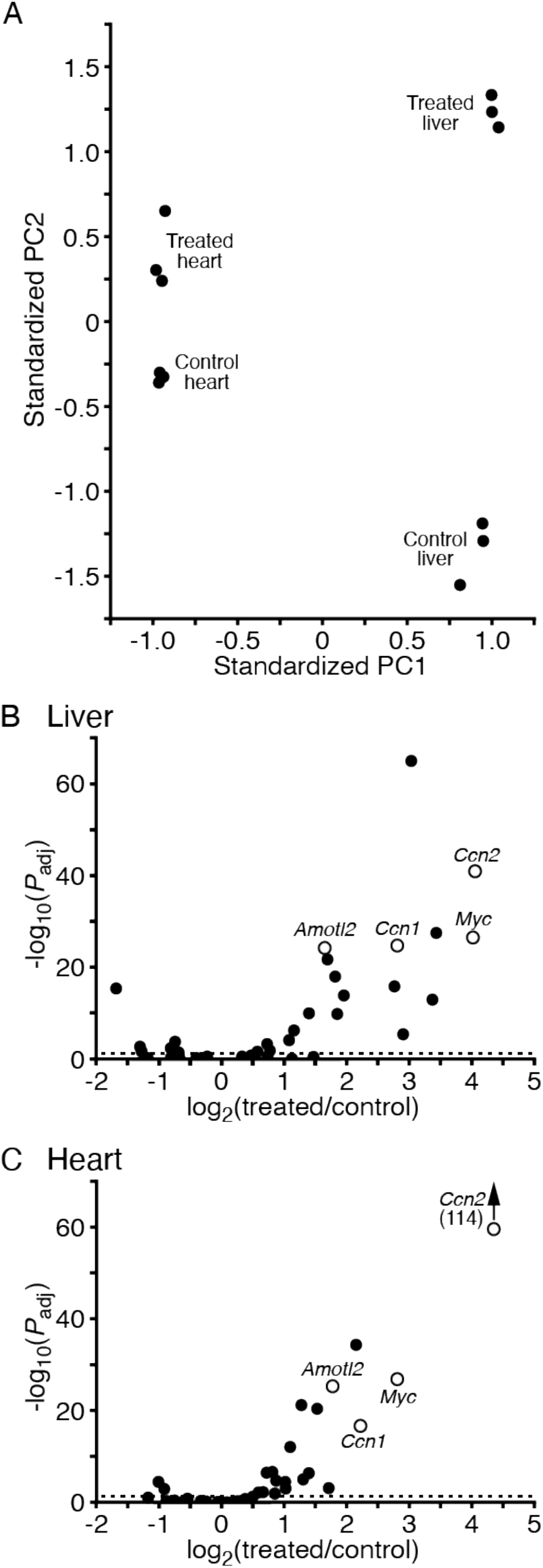
The effects of TDI-011536 treatment on gene expression. (*A*) Principal-components analysis of gene expression in the liver and heart indicates that the data are separated first on the basis of the respective organs, then in response to treatment with TDI-011536. The first principal component (PC1) captures 87.5 % of the explained variance and the second (PC2) accounts for 3.8 %. (*B*) A volcano plot demonstrates upregulation of Yap target genes after exposure to TDI-011536 in the liver. Examples of well-characterized targets of Yap are indicated as hollow circles: *angiomotin-like 2* (*Amotl2*), *cellular communication network factor 1* (*Ccn1*), *cellular communication network factor 2* (*Ccn2*), and *myc proto-oncogene* (*Myc*) The dashed horizontal line represents an adjusted probability level of *P*_adj_ = 0.05. (*C*) The corresponding volcano plot for the heart reveals a similarly robust enhancement of expression of Yap target genes. All data are available through GEO (submission 196322).

To examine potential toxicity in these tissues, we used gene set-enrichment analysis to inquire whether the gene ontology terms associated with inflammation (GO:0006954) or apoptosis (GO:0006915) were enriched after treatment. Although in both tissues we observed an enhancement of genes associated with inflammation, we could not demonstrate enrichment of any daughter gene ontology terms; the effect might therefore be non-specific. Although some genes associated with apoptosis were upregulated, the entire group did not display significant enrichment (SI Appendix, Fig. S2B). Further gene-ontology analysis of the genes enriched or reduced in both categories failed to identify a clear directionality of the effect. These results suggest that four hours’ treatment *in vivo* with TDI-011536 did not elicit significant damage in the organs examined.

### *In vivo* proliferative effect of TDI-011536 on cardiomyocytes

Cardiomyocytes ordinarily display little or no proliferation in mature mammals; instead, cardiac lesions in mice evoke scarring (12). To ascertain whether TDI-011536 restores a capacity for proliferation, we therefore examined the effect of systemic administration of the compound on damaged hearts in mice. After optimizing the surgical approach and to establishing an appropriate dosage regime, we conducted eight experiments on animals eight weeks of age. In each instance, we used a metal probe cooled in liquid nitrogen to create a small cryolesion on the right ventricle (12). We then inquired whether the cardiomyocytes near the affected area demonstrated proliferation, as marked by the incorporation of EdU.

Two or three days’ intraperitoneal dosing of six animals with 100 mg/mL of TDI-011536 resulted in the survival of all the mice for two weeks with no apparent ill effects. Two control animals were lesioned and injected with vehicle on the identical schedule. Outside the actual lesions, where numerous cells proliferated in the course of scar formation, EdU-positive cells were rare in the controls and few if any labeled cardiomyocytes were observed (Fig. 5A). In treated animals, on the other hand, EdU-labeled cells were widely scattered throughout the myocardium (Fig. 5B). Examination of the tissue at high magnification revealed numerous labeled cardiomyocytes (Fig. 5C-J).

**Figure 5.**
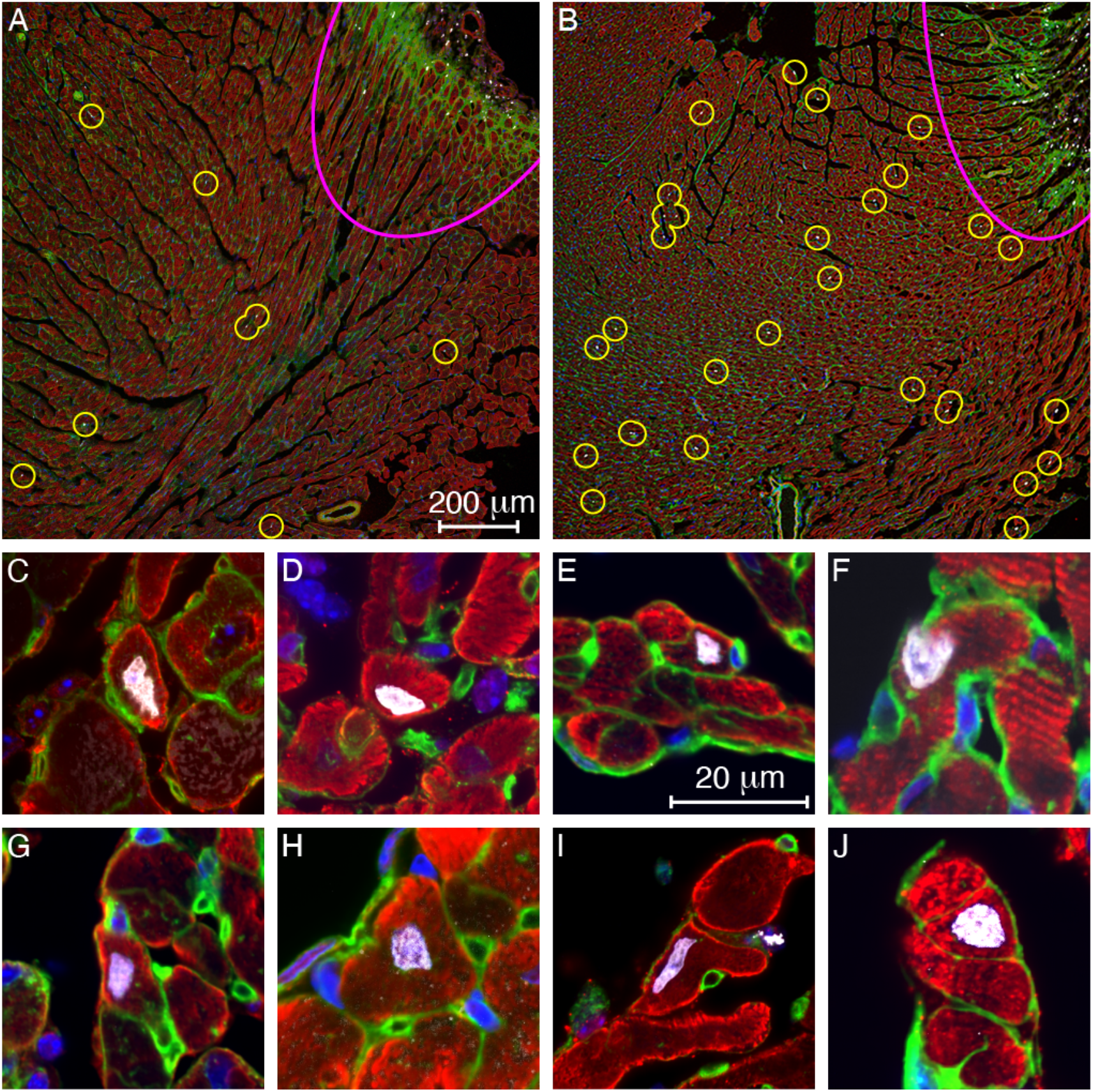
EdU incorporation into cardiomyocytes treated with TDI-011536. (*A*) At a distance from the lesion (pink arc at top right), a low-power micrograph of a representative section from a control animal shows only a handful of small EdU-positive cells (circles). (*B*) After three days’ treatment with TDI-011536, there are substantially more EdU-labeled cells outside the lesion, many of them large cardiomyocytes. Panels (*C*) through (*J*) display individual EdU-labeled cardiomyocytes from treated animals. In all the images, cardiomyocytes are immunolabeled for troponin I or alpha-smooth muscle actin (red). Nuclei are stained with DAPI (blue) and those incorporating DNA precursors are additionally marked with EdU (white). By labeling membrane glycoproteins, wheat-germ agglutinin (green) helps to delineate the boundaries of cardiomyocytes. The small round profiles represent capillaries.

In 33 images from 13 sections from the hearts of the two animals treated with TDI-011536 for three days, the average density of EdU labeling was 17.0 ± 3.5 µm^-2^ (mean ± SD; SI Appendix, Table S1). By contrast, 28 images from 12 sections from controls yielded a labeling density of 5.6 ± 2.1 µm^-2^ (mean ± SD; SI Appendix, Table S2). Because the values for the two control samples were not statistically different, those data were combined. Both treated samples showed significantly more labeling than controls, one with *P* < 10^−9^ and the other with *P* < 10^−14^ by one-tailed *t*-tests.

## Discussion

Because of its ubiquitous role in suppressing cell division, the Hippo-Yap phosphorylation cascade represents a potential impediment to regenerative therapies. Although the pathway’s activity might be modulated at several upstream levels, the key downstream targets are Lats1 and Lats2, paralogous protein kinases that directly modulate the activity of the transcriptional co-activators Yap and Taz. When unphosphorylated, Yap and Taz accumulate in the nucleus and enhance the expression of a variety of proteins associated with the cell cycle and mitosis. The small molecule TRULI, which blocks both Lats kinases, has therefore proven effective *in vitro* at triggering proliferation in mammalian organs as recalcitrant as the inner ear, retina, and heart (11).

In the present study, we have examined analogs of TRULI in an effort to determine which aspects of the molecule can be altered to enhance properties such as Lats inhibition, drug-like properties, and *in vivo* efficacy. A promising derivative, TDI-011536, demonstrates significant improvements in both potency and stability. Of equal importance, upon systemic administration the compound is not unduly toxic, reduces the phosphorylation of Yap for several hours in multiple organs *in vivo*, and evokes a modest degree of DNA synthesis in cardiomyocytes after focal cardiac damage.

Murine cardiomyocytes generally lose the capacity for proliferation by the first week after birth, but a burst of proliferation follows a week later owing to thyroid stimulation and some regenerative capacity persists for a few additional weeks (13)(14)(15). Although the majority of ventricular cardiomyocytes in mature mice are binucleated (16), the expression pattern of genes related to proliferation suggests that the minority population of mononucleated cells retains a greater proliferative capacity (17). Our results do not indicate the nuclear status of the cells that incorporate EdU and might therefore contribute to regeneration of damaged heart tissue. Moreover, we have not yet investigated the fate of the EdU-positive cells, which might undergo polyploidy or polynucleation without cytokinesis (13).

Although compounds related to TDI-011536 might have utility in regenerative therapies, accomplishing this would require overcoming several potential obstacles. In many instances the suppression of Lats activity alone is unlikely to suffice for a proliferative effect. In experiments on the internal ear *in vitro*, for example, TRULI provokes robust mitosis in supporting cells of a vestibular organ, the utricle, yet fails to elicit such a response in the auditory organ of Corti (11). Even though compounds of this class might release one brake on cellular proliferation, there are doubtlessly others that would need to be independently inactivated.

Potential toxicity is a second concern. The human proteome contains over 500 protein kinases (18), many with similar binding sites for ATP. As a result, it is extraordinarily difficult to target a particular kinase without off-target effects. TRULI, the progenitor of TDI-011536, blocks several off-target kinases, but few of major importance (11). To realize any therapeutic potential of compounds of this class, it would be necessary to profile in detail their interaction with a wider variety of kinases and to conduct further medicinal chemistry in an effort to reduce side effects.

A related issue is the mechanism of dosage. Systemic administration of a compound such as TDI-011536 obviously risks deleterious effects in numerous organs, and indeed the present study was restricted in young mice by lethality related to intraperitoneal injection. However, in many instances it should be possible to limit this problem by focused administration of a compound, for example into the perilymph of the internal ear, the vitreous body of the eye, or the pericardium of the heart. Localized administration by a mini-pump or other apparatus for focal infusion might prove effective as well.

The duration of exposure is also an important consideration. Any treatment that suppresses the normal restrictions on cellular proliferation could prove dangerous as a result of unwanted hyperplasia or even carcinogenesis. An attractive feature of treatment with a substance related to TDI-011536, however, is that such compounds evoke proliferation within only a few days (11) and need not be administered for any longer period: in most instances, one or a few rounds of mitosis could restore a damaged organ’s function. Moreover, although some hyperplasia was observed during treatment with a compound that blocks upstream components of the Hippo-Yap pathway, the Mst kinases, the affected organs reverted to normal size within days after exposure without an increased incidence of cancer during the subsequent year (8).

Two other characteristics of potential therapeutic agents could be improved by additional chemical intervention. It is likely that the efficacy of the compounds could be further enhanced: some inhibitors already in hand block Lats kinases at sub-nanomolar concentrations even in the presence of 2 mM ATP and suppress pYap with an *EC*_50_ in the low nanomolar range in HEK293A cells. Although greater potency might be associated with more off-target activity, there are likely compounds both more potent and more selective than TDI-011536. A second feature that could be improved is solubility. With few polar groups, TRULI and TDI-011536 are sparingly soluble in aqueous media. As further investigations—especially crystallography— clarify the interactions of such compounds with Lats kinases, it should become apparent at what sites polar groups might be added to enhance solubility and thus to facilitate the administration of potential drugs.

## Materials and Methods

All procedures involving living animals were conducted with the approval of Rockefeller University’s Institutional Animal Care and Use Committees.

Data relating to gene expression in response to TDI-011536 are available at GEO (submission GSE196322; https://www.ncbi.nlm.nih.gov/geo/query/acc.cgi?acc=GSE196322).

### Assays of Lats kinase inhibition

The activity of compounds was tested against a functional component of Lats kinases in an enzymological assay (11). The substances’ ability to reduce the phosphorylation of Yap was also tested in HEK 293 cells, again as described previously. For the latter assay, variations in the control responses sometimes led to negative values or values exceeding 100 % for the percentage of inhibition, but the dependence of the results on the concentrations of the test compounds was highly reproducible.

### Systemic administration of TDI-011536

The relevant compound was emulsified by sonication in 10 % Kolliphor HS 15 (42966, Sigma) in phosphate-buffered saline solution (10010-023, Gibco). The mixture was administered intraperitoneally at 50-200 mg/kg of TDI-011536. When appropriate, EdU (E10187, Thermo Fisher) was administered intraperitoneally at a dosage of 5 mg/kg on each day of treatment.

### Measurement of *in vivo* Lats inhibition

Fifty milligrams of heart, liver, or skin tissue was lysed on ice in radioimmunoprecipitation-assay (RIPA) buffer solution (BP-115, Boston BioProducts) with protease inhibitors (Halt Protease Inhibitor Cocktail 87786, Thermo Fisher). After remaining on ice for 30 min, the samples were sonicated at low power for 2 min with 10 s exposures separated by 10 s interruptions, then centrifuged at 21,130 X g for 10 min at 4 °C. The supernatants were filtered and the lysates were immediately subjected to electrophoresis or stored at -80 °C.

A standard immunoblotting protocol was used. After lysates had been prepared, 10 mg of protein from the heart or liver or 20 mg of protein from the skin was resolved on a 4 %-12 % gradient bis-tris gel (NP0322, Thermo Fisher). The proteins were transferred to a PVDF membrane (1704156, BioRad) and blocked for one hour at room temperature (MB-070, Rockland). We used primary antibodies directed against Yap (101199, Santa Cruz Biotechnology), phosphorylated Yap S127 (4911, Cell Signaling Technology) and Gapdh (ab8245, Abcam), which were reconstituted at a dilution of 1:1000 in blocking solution. After a membrane had been exposed to primary antibodies overnight at 4 °C, it was subjected to three 5 min washes in tris-buffered saline solution (28358, Thermo Fisher) with 0.05 % Tween-20. Secondary antibodies conjugated to horseradish peroxidase (1:10,000, W401B and W402B, Promega) were applied in the same solution for one hour at room temperature before activity was detected (SuperSignal West Pico PLUS 34580, Thermo Fisher). Images were acquired with an iBright FL1000 system (Invitrogen).

### Culture of human pluripotent stem cells and retinal organoids

After induced pluripotent stem cells of the WTC-11 line (Coriell Institute for Medical Research, Camden, NJ) had been grown by standard methods, human retinal organoids were produced (19). The effects of Lats kinase inhibition were analyzed as described previously (11). For the preparation of protein lysates after a 24-hour treatment in two independent assays, five organoids 225-280 days of age were sampled per experimental group. In four independent proliferation assays, the organoids were incubated in long-term culture medium supplemented with 10 µM EdU and 10 µM TRULI or 3 µM TDI-011536. The extent of proliferation was assessed by quantifying the percentage cells doubly positive for EdU and Sox9 after five days in culture.

### RNA sequencing

Sequence and transcript coordinates for the murine mm10 UCSC genome and gene models were retrieved from the Bioconductor Bsgenome.Mmusculus.UCSC.mm10 (version 1.4.0) and TxDb.Mmusculus.UCSC.mm10.knownGene (version 3.4.0) Bioconductor libraries, respectively. Transcript expression was calculated with Salmon quantification software (version 0.8.2) (20); gene expression levels as transcripts per million and counts were retrieved with Tximport (version 1.8.0) (21). Normalization and rlog transformation of raw read counts in genes were performed with DESeq2 (version 1.20.0) (21). *P*-values were corrected for multiple testing by the Benjamini-Hochberg algorithm. Intersample variability was assessed with hierarchical clustering, and heat maps of intersample distances were constructed in the Pheatmap R package (version 1.0.10). Gene set-enrichment analysis was conducted through clusterProfiler (version 3.18.1) (22). Other plotting was performed using ggplot2 (version 3.3.3). Potential targets of Yap signaling were identified from studies of breast cancer cells (23).

### Cryolesioning of the murine right ventricle

Cardiac lesions were produced by a variant of a published method (12). Twenty minutes before surgery, an adult Swiss Webster mouse (Charles River Laboratories) four or eight weeks of age and of either sex was given buprenorphine subcutaneously at a dosage of 100 µg/kg for analgesia. The animal was anesthetized with 100 mg/kg of ketamine and 5 mg/kg of xylazine administered intraperitoneally in sterile water. The animal’s mid-ventrum was shaved, chemically depilitated, and sterilized with a povidine-iodine swab. The area of the skin incision was infiltrated intradermally with 1 mg/kg of bupivacaine. Subsequent procedures were conducted beneath a surgical microscope and under sterile conditions in a HEPA-filtered, positive-pressure hood.

With the animal supine on a thermostatted heating pad at 40 °C, an oblique, 15 mm-long incision was made from the xiphoid process parallel to the left lower costal margin and about 3 mm caudal to it. After the abdominal musculature and peritoneum had been transected with scissors in the same pattern, a miniature retractor was introduced to separate the two sides of the incision. This approach exposed the upper margin of the liver and provided a view of the beating heart and the base of the left lung through the transparent diaphragm.

A cryolesion of the right ventricle was created with a round aluminum rod of mass 1.44 g secured to a plastic handle. A total of 76 mm in length, the rod was 3.2 mm in diameter over a distance of 59 mm at the base and 2.15 mm in diameter over the 17 mm at the tip. The entire rod was cooled for 20 s by immersion in liquid nitrogen, after which the slightly rounded, polished tip was pressed against the diaphragm near the middle of the roughly rectangular area of exposure of the heart. A constant force, estimated as 0.8 N by simulation with an electronic balance, was exerted for 20 s, after which the probe was withdrawn. A successful lesion was marked by a 2 mm disk of frozen diaphragm—which rapidly thawed—and often by a dark patch on the subjacent heart.

It proved important to avoid two possible complications. First, it was necessary to insert the probe at a steep enough angle with respect to the horizontal to avoid touching and lesioning the liver. And second, it was essential not to exert excessive force: if the diaphragm wrapped around the end of the probe, it adhered so strongly that its withdrawal occasionally perforated the diaphragm and caused an immediately fatal pneumothorax. After preliminary experiments, however, all of the five to ten mice operated on a given day survived the procedure and became active in a warmed cage 10-30 min after surgery. To maintain analgesia, each animal was administered an additional 100 µg/kg of buprenorphine subcutaneously at 12-hour intervals for each of the days on which TDI-011536 was injected.

### Histological preparation and immunolabeling of cardiac muscle

After an animal had been anesthetized and sacrificed by cervical dislocation, its heart was rapidly removed. The site of the lesion was apparent as a 2 mm purple spot in the middle portion of the right ventricle (12). To facilitate the access of fixative, the base and apex of the heart were amputated and the lesioned segment was immersed for 16 hours in 4 % formaldehyde in phosphate-buffered saline solution (28906, Thermo Fisher). The specimen was then immersed overnight at 4 °C in 30 % sucrose solution, then infiltrated with cryoprotectant (OCT), frozen, and sectioned at a thickness of 5-8 µm.

Mounted on a glass slide, each section was blocked in a humidified chamber for one hour at room temperature with 3 % bovine serum albumin (AB 2336846, Jackson), 5 % normal donkey serum (AB-2337258, Jackson), and 0.5 % Triton X-100 (93443, Sigma) in tris-buffered saline solution. For the detection of EdU, the sections were then subjected to click chemistry (Click-iT EdU imaging kit C10340, Thermo Fisher) for 30 min at room temperature. After a short rinse with phosphate-buffered saline solution, the sections were labeled overnight at 4 °C with Alexa Fluor 488-labeled wheatgerm agglutinin (1:200; W11261, Thermo Fisher) in combination with a rabbit polyclonal primary antiserum, either anti-cardiac troponin I (1:50; ab47003, Abcam) or anti-alpha smooth muscle actin (1:50; ab5694, Abcam). After slides has been washed three times with phosphate-buffered saline solution, Alexa-Fluor 555-labeled secondary antiserum (1:500; A32794, Invitrogen) diluted in blocking solution was added for one hour at room temperature followed by two washes with phosphate-buffered saline solution. Nuclei were stained with DAPI and sections were mounted with glass cover slips in a mounting medium (Prolong Gold Anti-fade P36934, Thermo Fisher).

Low-power images were obtained with a confocal microscope (LSM 780, Zeiss) equipped with a 10X plan-apochromatic objective lens of numerical aperture 0.45 The images were processed with Fiji (24) to estimate the density of EdU-labeled cells (SI Appendix, Fig. S3). High-power images were acquired on an inverted microscope (IX81, Olympus) equipped with a super-resolution fluorescence illumination system (VT-iSIM, VisiTech International) and a 60X silicone-oil-immersion objective lens of numerical aperture 1.3 (UPLSAPO60XS2, Olympus).

## Acknowledgments

The authors thank Dr. Peter Meinke for consultation and directorship of the Tri-Institutional Therapeutics Discovery Institute, a 501(c)(3) organization that receives funding from its parent institutes—The Rockefeller University, Memorial Sloan Kettering Cancer Center, and Weill Cornell Medical College—and from Takeda Pharmaceutical Company; from Mr. Lewis Sanders; and from other philanthropic sources. Dr. Leslie Diaz of the Comparative Bioscience Center provided expert advice about surgical procedures. Some micrographs were obtained at the Bio-Imaging Resource Center, and the inhibition of Lats kinases in cells was conducted at the High-Throughput Screening Resource Center, both at The Rockefeller University. Retinal organoids were produced at the Stem Cell Analytics Core and imaged at the Cellular Imaging Core, both at The Saban Research Institute of Children’s Hospital Los Angeles. We thank Eva Jahanshir, Sravani Ramisetty, and Kayla Stepanian for technical assistance with organoid culture and analysis. We also acknowledge helpful comments about the manuscript from our research groups. N.K. was supported by the NIGMS (T32GM007739); A.N. and K.G. are supported by the Edward N. & Della L. Thome Memorial Foundation Awards Program in Age-Related Macular Degeneration Research; A.N. is also supported by a Research to Prevent Blindness Career Development Award, The Leon and Carol Ellison Research Career Development Award at CHLA, the Las Madrinas Endowment in Experimental Therapeutics for Ophthalmology, and the NEI (K08EY030924); and K.G. by the NIDCD (R21DC016984). A.J.H. is an Investigator of Howard Hughes Medical Institute.

## Table

**Table 1.** Upregulation of Yap-target genes by TDI-011536.

| Gene | Heart — — | — — — — — | — — — — — | Liver — — | — — — — — | — — — — — |
| --- | --- | --- | --- | --- | --- | --- |
|  | Control | Treated | Significance | Control | Treated | Significance |
| *Amotl2* | 9.77 ± 0.09 | 11.53 ± 0.13 | 5·10<sup>-26</sup> | 8.58 ± 0.05 | 10.21 ± 0.07 | 5·10<sup>-25</sup> |
| *Ccn1* | 9.28 ± 0.16 | 11.46 ± 0.29 | 2·10<sup>-17</sup> | 6.77 ± 0.06 | 9.50 ± 0.27 | 2·10<sup>-25</sup> |
| *Ccn2* | 9.66 ± 0.18 | 14.02 ± 0.14 | 9·10<sup>-115</sup> | 6.59 ± 0.14 | 10.55 ± 0.33 | 1·10<sup>-41</sup> |
| *Myc* | 5.42 ± 0.18 | 8.15 ± 0.10 | 1·10<sup>-27</sup> | 7.23 ± 0.32 | 10.67 ± 0.23 | 3·10<sup>-28</sup> |

Each measurement shown is an rLog statistic that represents a modified log<sub>2</sub> transformation regularized by variance stabilization. This approach accounts for the relationship between variance and mean in RNA-seq data by shrinking the observed variance for genes with low counts. By preventing the data from genes with low counts from becoming spread apart after logarithmic transformations, the procedure allows meaningful signals to better emerge from the background noise. Uncertainties are shown as SEMs for three measurements in each instance. Significance values were determined by two-tailed Wald tests.

## Supporting Information figures

**Figure S1.**
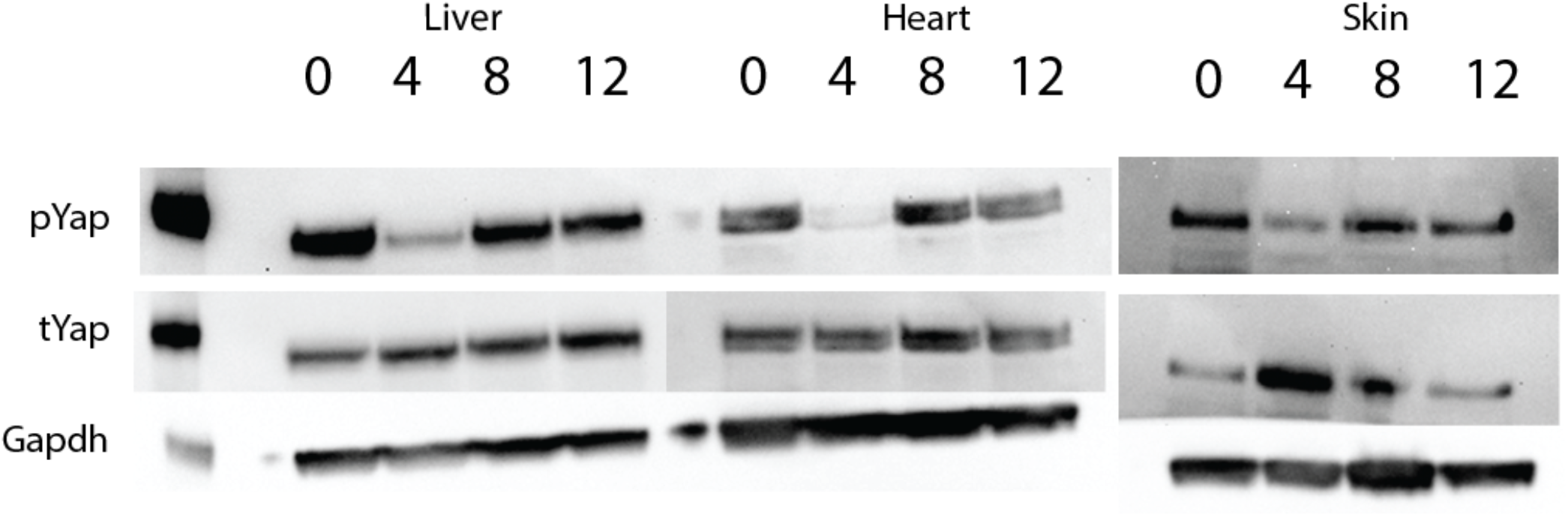
Recovery of pYap levels. Immunoblots portray the amounts of Yap phosphorylated at residue S127 (pYap) and the total amounts of Yap (tYap) in the liver, heart, and skin. Although intraperitoneal administration of TDI-011536 at a dosage of 200 mg/kg of body weight reduced the level of phosphorylation by four hours after injection, recovery was essentially complete by eight hours in all three organs. Glyceraldehyde 3-phosphate dehydrogenase (Gapdh) is included in each instance as a loading control.

**Figure S2.**
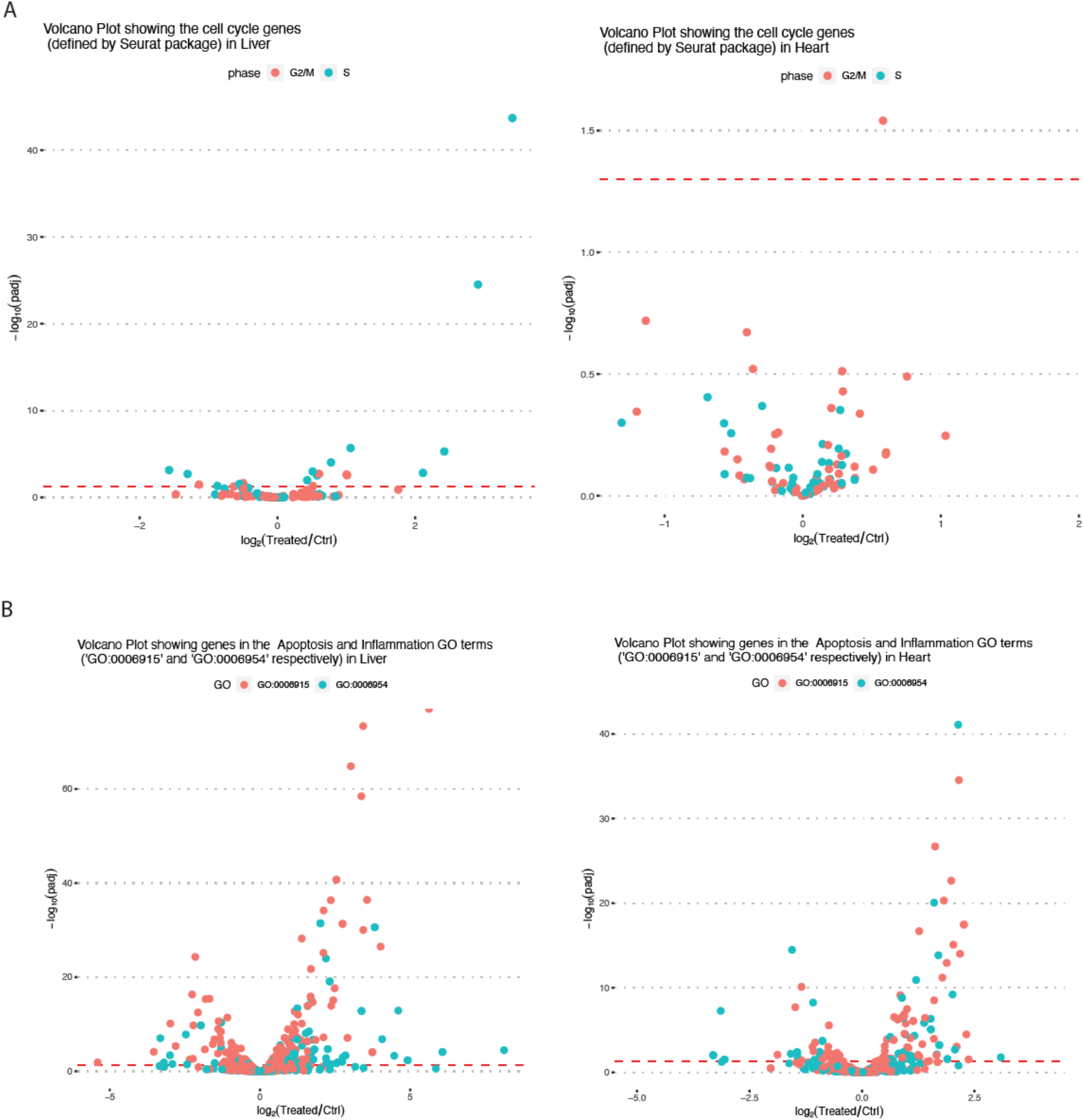
Effect of TDI-011536 on expression of genes in liver and heart associated with mitosis, inflammation, and apoptosis. These volcano plots demonstrate the individual genes from the respective GO term sets, with their fold changes presented as log_2_(treated/control) on the abscissa and the significance plotted on the ordinate as -log_10_(adjusted *P*-value). (*A*) Four hours after intraperitoneal administration of TDI-011536 at a dosage of 200 mg/kg in liver and heart, a small number of genes from the G2/M (red) and S (teal) GO terms are significantly enriched, but the group as a whole is not significantly up-regulated by gene set-enrichment analysis. (*B*) After the same treatment, the Inflammation GO set (teal) is significantly up-regulated in both liver and heart. Some of the Apoptosis-associated genes (red) are significantly up-regulated, but gene set-enrichment analysis does not find the group as a whole enriched. Further gene-ontology analysis of the genes enriched or reduced in both categories fails to identify a clear directionality of the effects. Dotted red lines indicate adjusted *P*-values of 0.05.

**Figure S3.**
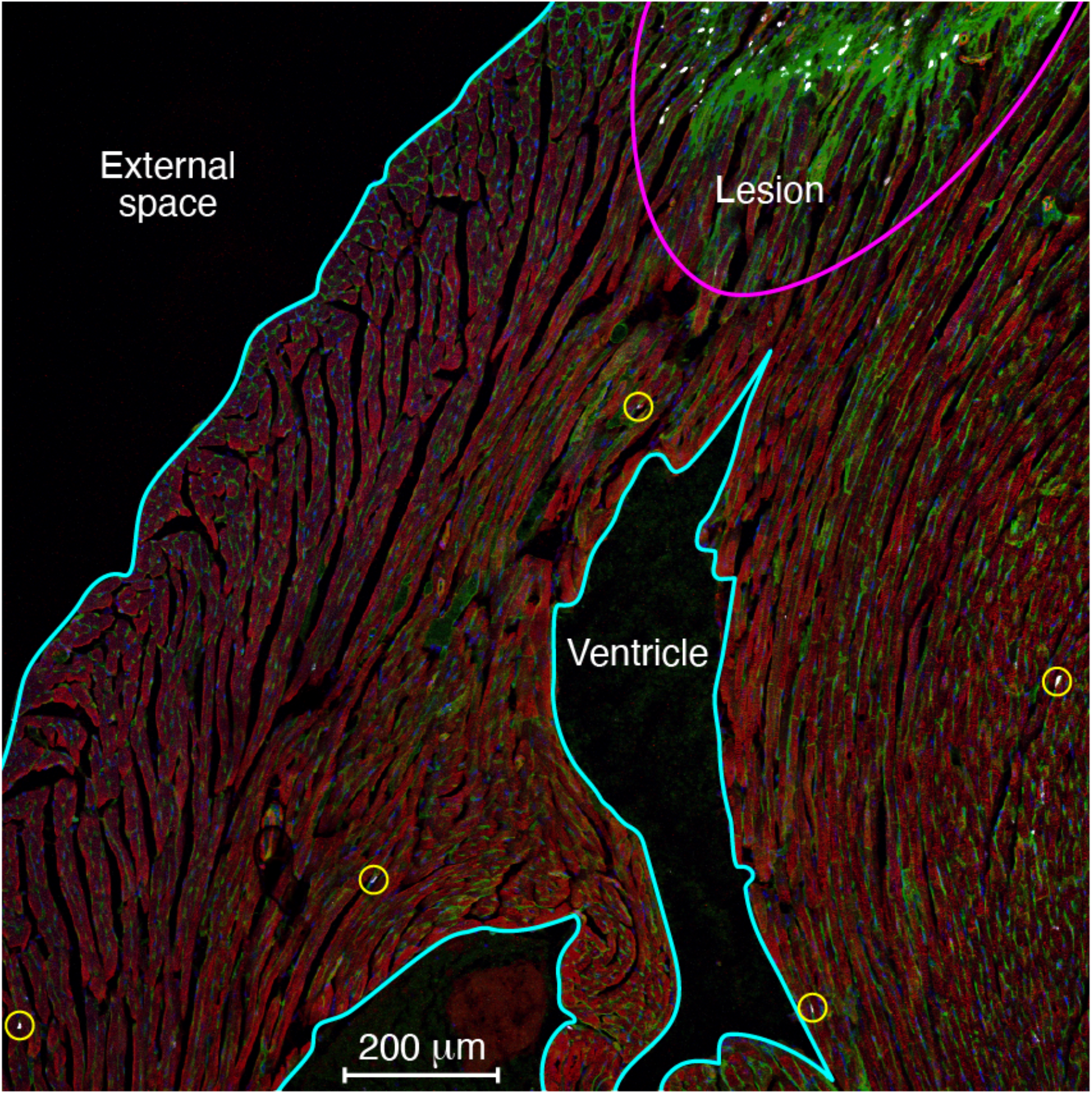
Procedure for estimation of Edu labeling density. In a low-power micrograph of right ventricular tissue from a control animal, a cryolesion (pink arc) is marked by enhanced labeling with wheat-germ agglutinin (green) owing to the elimination of the cells formerly in that area. Other areas free of muscle are outlined (light blue). The combined area of the lesion, the ventricular cavities, and the external space is 0.631 mm^2^, or 31.4 % of the total image area of 2.008 mm^2^. The 68.6 % of the image containing cardiac muscle and lying outside the lesion— an area of 1.377 mm^2^—contains five EdU-positive cells (yellow circles); the labeling density is accordingly 3.63 mm^-2^.

## Supporting Information tables

**Table S1.** Density of EdU-labeled cells in TDI-011536-treated preparations.

| Animal label | Slide label | Image number | Nonmuscle area (mm<sup>2</sup>) | Muscle area (mm<sup>2</sup>) | Labeled cell count | Density (mm<sup>-2</sup>) |
| --- | --- | --- | --- | --- | --- | --- |
| Specimen 1 | AB | 1.1 | 0.239 | 1.769 | 24 | 13.57 |
|  |  | 1.2 | 0.216 | 1.792 | 30 | 16.74 |
|  |  | 1.4 | 0.810 | 1.198 | 25 | 20.87 |
|  |  | 3.1 | 0.620 | 1.388 | 23 | 16.57 |
|  | AC | 1.1 | 0.127 | 1.881 | 28 | 14.89 |
|  | AG | 1.2 | 0.341 | 1.667 | 21 | 12.60 |
|  |  | 2.2 | 1.049 | 0.959 | 15 | 15.64 |
|  | AK | 1.1 | 0.443 | 1.565 | 18 | 11.50 |
|  | M | 1.2 | 0.525 | 1.483 | 19 | 12.81 |
|  |  | 2.1 | 0.223 | 1.785 | 24 | 13.45 |
|  |  | 3.1 | 0.497 | 1.511 | 23 | 15.22 |
|  | S | 3.1 | 0.990 | 1.018 | 22 | 21.61 |
|  | AV | 1.1 | 0.544 | 1.464 | 21 | 14.34 |
|  |  | 3.1 | 0.677 | 1.331 | 18 | 13.52 |
| Specimen 2 | AB | 2.1 | 0.809 | 1.199 | 18 | 15.01 |
|  |  | 2.2 | 0.104 | 1.904 | 34 | 17.86 |
|  | AG | 1.1 | 0.435 | 1.573 | 26 | 16.53 |
|  |  | 1.2 | 0.519 | 1.489 | 21 | 14.10 |
|  |  | 2.1 | 0.409 | 1.599 | 27 | 16.89 |
|  | K | 1.1 | 0.629 | 1.379 | 21 | 15.23 |
|  |  | 1.2 | 0.318 | 1.690 | 27 | 15.98 |
|  |  | 2.1 | 0.444 | 1.564 | 32 | 20.46 |
|  |  | 2.2 | 0.301 | 1.707 | 34 | 19.92 |
|  |  | 3.1 | 0.250 | 1.758 | 37 | 21.05 |
|  |  | 3.2 | 0.204 | 1.804 | 28 | 15.52 |
|  | P | 1.1 | 0.365 | 1.643 | 41 | 24.95 |
|  |  | 1.2 | 0.227 | 1.781 | 35 | 19.65 |
|  | S | 1.1 | 0.443 | 1.565 | 24 | 15.34 |
|  |  | 1.2 | 0.730 | 1.278 | 20 | 15.65 |
|  |  | 2.1 | 0.163 | 1.845 | 41 | 22.22 |
|  |  | 2.2 | 0.220 | 1.788 | 38 | 21.25 |
|  |  | 3.1 | 0.190 | 1.818 | 29 | 15.95 |
|  |  | 3.2 | 0.365 | 1.643 | 41 | 24.95 |

Each image captured a field of $1417\ \mu\text{m} \times 1417\ \mu\text{m}$ and therefore a total area of $2.008\ \text{mm}^2$ . The density of labeled cells in 14 images from Specimen 1 was $15.24 \pm 2.95\ \text{mm}^{-2}$ and that in 19 images from Specimen 2 was $18.34 \pm 3.39\ \text{mm}^{-2}$ (means ± SDs). The two results were significantly different at the level $P = 0.0088$ by a double-tailed $t$ -test with unequal variances. Combining the data yielded a density of $17.0 \pm 3.5\ \text{mm}^{-2}$ (mean ± SD). A comparison of these combined data with the combined control data showed enhanced labeling of treated specimens at the level $P < 10^{-21}$ by a single-tailed $t$ -test with unequal variances.

**Table S2.**
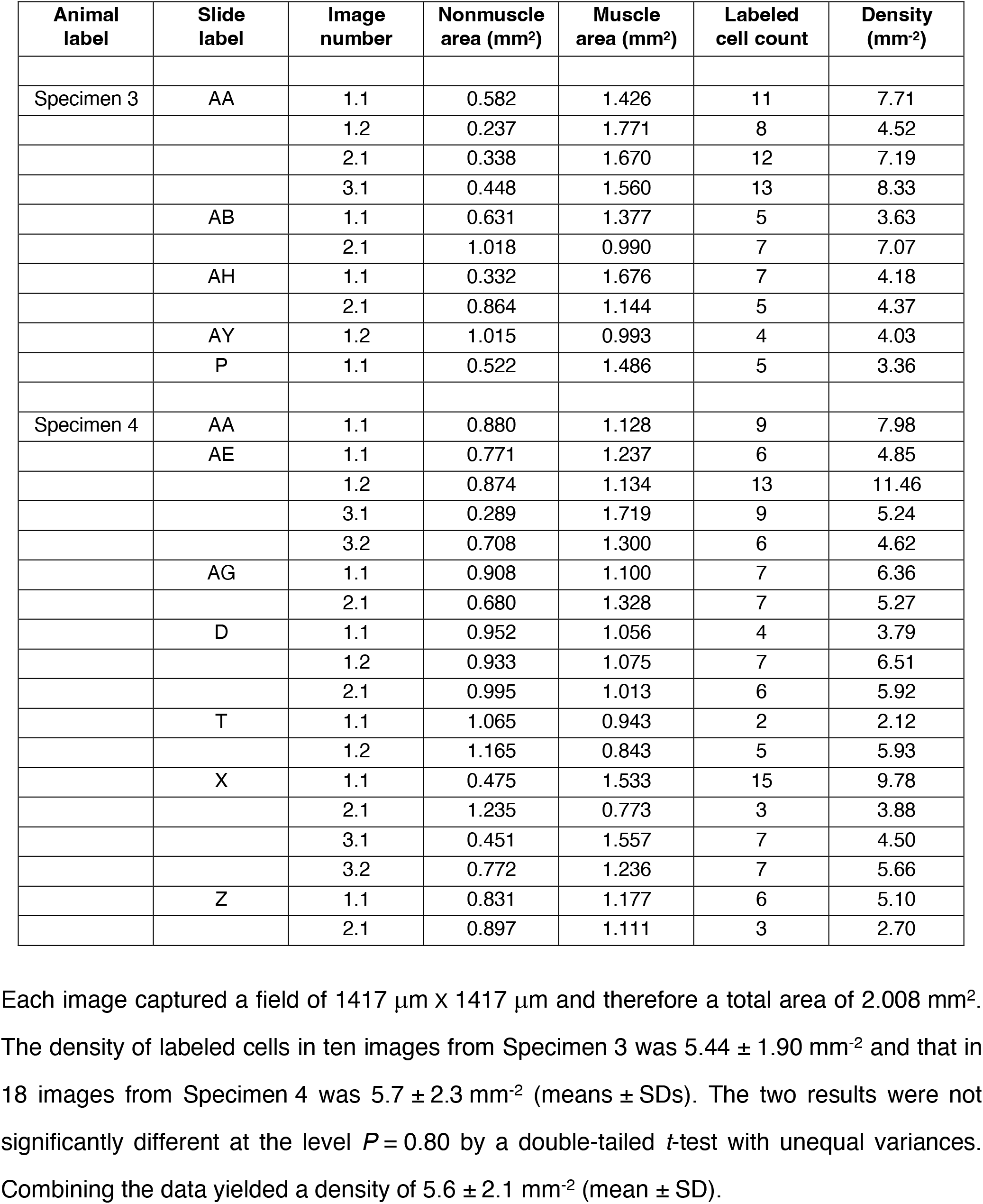
Density of EdU-labeled cells in control preparations.

